# Demultiplexing overlapping signaling scaffold functions to probe lipid messenger coupling to cytoskeletal dynamics

**DOI:** 10.1101/657049

**Authors:** Nicholaus J. Trenton, R. Tyler McLaughlin, Satya K. Bellamkonda, David S. Tsao, Emily M. Mace, Jordan S. Orange, Volker Schweikhard, Michael R. Diehl

**Affiliations:** Department of Bioengineering, Rice University, Houston, TX 77030, USA; Graduate Program in Systems, Synthetic and Physical Biology, Rice University, Houston, TX 77030, USA; Department of Pediatrics, Columbia University Irving Medical Center, New York, New York; Department of Chemistry, Rice University, Houston, TX 77030, USA

## Abstract

The coordination of lipid messenger signaling with cytoskeletal regulation is central to many organelle-specific signaling and regulatory processes. While central to many aspects of cell physiology, this coupling often depends on the function of multi-domain scaffolds that orchestrate transient interactions and dynamic feedback among a spectrum of signaling intermediates and regulatory proteins on organelles. Understanding scaffold protein functions has remained challenging given this complexity. This work employs live-cell imaging and statistical analyses to deconvolve (demultiplex) how the multi-domain scaffold IQGAP1 coordinates phosphoinositide signaling with organelle-specific actin regulation and membrane processing events. Using actin-ensconced endosomes that localize to the basal cortex of polarized epithelial cells as a model system, we demonstrate abilities to dissect how IQGAP1 transitions between different actin and endosomal-membrane tethered states. We provide evidence IQGAP1 functions as a transient inhibitor of actin growth around the endosomes in at least one of these states. While not easily distilled via standard (static) colocalization analyses or traditional pathway perturbations methods, this negative regulation was revealed via a series of dynamic correlation and multiple regression analyses. These methods also uncovered that the negative actin regulation is linked to GTPase-dependent tethering to the endosomal membrane. Moreover, the scaffold transitions underlying this control are shown to depend on the production of PIP_3_ lipid messengers by the lipid kinase PI3K. Overall, these methods and results provide new insights in to how IQGAP1 act as a signaling hub by orchestrating time-dependent membrane and cytoskeletal protein interactions and provide new routes to dissect scaffold-mediated pathway regulation in a variety of settings.

## INTRODUCTION

Phosphoinositides (PIs) are an important class of lipid messenger that drive the recruitment of cytoplasmic signaling proteins to the plasma membrane, organelles, and vesicles (1). PIs also participate in signal transduction by activating proteins locally and relaying signaling events between membrane receptors (2), ion channels (3), as well as downstream membrane and cytoskeletal regulatory proteins (4-6). This communication is at the center of mechanisms governing organelle development, and cell growth, polarization, death, proliferation and motility. Aberrant PI signaling has been associated with numerous human diseases (7). For example, multiple oncogenic mutations have been shown to cause hyperactivation of phosphatidylinositol (3,4,5)-trisphosphate (PIP_3_) dependent signaling in the phosphoinositide 3-kinase (PI3K) / Akt pathway at the early stages of many common cancers (8). Unfortunately, efforts to target PIP_3_ effector proteins are often confounded by unwanted side effects due to the extensive crosstalk and feedback that occurs within lipid signaling networks (9). While integral to mechanisms that coordinate signaling with membrane and cytoskeletal regulatory events, this complexity arises from dynamic interactions among the different PI species, their associated kinases and phosphatases, and a spectrum of GTPase-dependent signaling intermediates and regulatory complexes. Methods to characterize these multi-faceted interactions are therefore critical to both understanding the functional roles of PIs in cell physiology and managing them in human diseases.

PI signaling is naturally expected to depend on the integrated functions of large multi-domain signaling scaffolds (10). These multifunctional master-regulatory proteins operate as dynamic templates that control the assembly of multiple-protein complexes and tether them to local membrane and cytoskeletal structures (11). This control is foundational to PI signaling since the affinities of many PI binding proteins are often quite low (12). Multi-valent binding interactions are therefore important to stabilize PI recognition and membrane association. Yet, beyond tethering, scaffolds can enhance, insulate or terminate signaling processes by controlling the coupling sequence and strength of interactions between different enzymes and their effectors (13). Scaffolds can also define local feedback circuitry and control signaling pathways locally within specific subcellular compartments (14). These high-order functions are increasingly recognized for their influence over the spatial location, amplitude and timing of PI signaling. Nevertheless, a number of challenges limit efforts to resolve how scaffolds configure PI signaling networks and couple them to downstream regulatory processes.

This work examines how the multi-domain scaffold IQGAP1 (*IQ motif containing GTPase activating protein 1*) helps coordinate PI3K / PIP_3_ signaling with actin and membrane dynamics within specific subcellular compartments, in this case sorting endosomes in polarized epithelial cells. IQGAP1 is a very large (∼190 kDa) multi-domain scaffold that is highly expressed in all human tissues (15) and known for its roles in multiple physiological processes including cell motility, endocytosis, phagocytosis, secretion, cytokinesis, and tumor cell invasion (16). It is particularly influential to epithelial integrity, polarization and compartmentalized signaling (17, 18). IQGAP1 scaffolds orchestrate the activities of multiple signaling intermediates and regulatory proteins via coordination of 6 key protein domains: an N-terminal calponin homology domain (CHD), a REPEATS region followed by a WW domain, four isoleucine and glutamine IQ motifs, a RasGAP-related domain (GRD) and a RasGAP C-terminal (RGCT) domain. This domain structure presumably allows IQGAP1 to anchor proteins to both PIP_2_-positive membranes and actin networks (*via the RGCT and CHD respectively*) (19). Although IQGAP1 does not function as a traditional GTPase activating protein (GAP), it can directly bind to the GTPases Cdc42 and Rac1 and has been shown to stabilize their GTP-bound form (*via the GRD*) (20). These interactions have been implicated in mechanisms that regulate actin polymerization. IQGAP1 has recently been reported to template the assembly of multiple PI kinase complexes and facilitate processive, multi-step PI signaling (21). Overall, multiple studies implicate IQGAP1 as a master regulator of processes requiring coupling between membrane receptors, lipid messenger and either membrane remodeling or cytoskeletal regulatory proteins. However, as with many scaffolding proteins, far fewer studies have explored how IQGAP1 coordinates the dynamic functions of its different domains in order to perform these high order functions.

Our group recently demonstrated that IQGAP1 participates in a local regulatory network that controls the assembly and disassembly of dynamic actin networks that confine sorting endosomes to the basal cortex of polarized epithelial cells (22). Live-cell analyses of dynamic IQGAP1 and actin intensities indicate IQGAP1 localization and actin regulation within these compartments are governed by multiple overlapping scaffold– compartment interactions. Moreover, we have shown that these interactions occur at differ timescales, yielding non-monotonic, frequency-dependent localization behaviors and scaffold / actin intensity correlations (22). For example, IQGAP1 and actin intensity tend to rise and fall together over long timescales (>10 minute) as compartments assembled and disassembled due to direct scaffold-actin binding. This yields positive intensity correlations in static images and at low temporal frequencies. Yet, IQGAP1 and actin correlate negatively at higher frequencies (∼ 1 min time scale). This behavior indicates transient IQGAP1 dissociation prompts the polymerization of endosome-specific actin filaments by relieving local actin inhibition temporarily. The actin networks in these compartments therefore appear to be remodeled via multiple reiterative cycles of scaffold disassociation, actin polymerization, and scaffold re-association and actin stabilization.

An integrated regulatory relationship involving a combination of positive low frequency and negative high frequency interactions may seem to deviate from traditional notions of how IQGAP1 helps control actin dynamics. The GRD and RGCT domains at the C-terminus of IQGAP1 are also widely reported to promote actin polymerization via their associations with Cdc42 or Rac1, and subsequent recruitment and activation of the Wiskott–Aldrich Syndrome protein (WASp) and the actin-related protein (Arp) 2/3 complex (23, 24). This type of positive actin regulation has been implicated in mechanisms that promote actin polymerization within lamellipodia and other cellular protrusions. Yet, these functions are not necessarily simple and straightforward. For example, imaging analyses have shown that IQGAP1 colocalizes with Arp2/3 at the protruding edges of certain types of migrating cells, but binds to retracting edge regions that have lost Arp3 signals in other cells (25). The latter anti-correlated IQGAP1 / Arp3 behavior is potentially indicative of a negative or inhibitory regulatory function. Conflicting positive and negative regulatory behaviors have also been found using IQGAP1 silencing via RNA interference, which has been shown to reduce actin-driven membrane ruffling in lamellipodia (24) but found to upregulate actin polymerization in other settings including the formation of the T-cell immunological synapse (26). Overall, as we propose for the endosome compartments found in epithelial cells, it is possible that IQGAP1 plays both inhibitory and supportive roles in actin polymerization in each of these examples. However, with different frequency-dependent responses, measured behaviors will depend on the experimental timescale and how underlying behaviors are sampled. Dissecting or ‘demultiplexing’ how scaffolds like IQGAP1 coordinate proteins within local cytoskeletal regulatory networks and, importantly, couple this regulation to lipid messenger and other intracellular signaling pathways ultimately requires new approaches that can account for mixtures of temporally distinct functions.

The present study further dissects the multifaceted dynamics of IQGAP1 and its role in actin regulation and phosphoinositide signaling via a series of multiplexed live-cell imaging analyses of wild type and mutant scaffold constructs and statistical modeling approaches. Using IQGAP1-positive sorting endosomes in MCF10A cells as a model compartment, we provide evidence that IQGAP1 controls the growth and disassembly of endosome-specific actin networks by transitioning between different actin and membrane bound states. The negatively-correlated, high frequency intensity fluctuations are further linked to IQGAP1-dependent inhibition of actin and shown to depend on the membrane tethering modes of the scaffold. Ties to membrane processing and tubule formation are also presented. These different activities were delineated via correlation and multiple regression analyses that employ different sets of scaffold mutants, actin labels and membrane markers. Importantly, multiple regression allows the mutant constructs to be used to deconvolve / ‘demultiplex’ the functions of scaffold domains. The IQGAP1 constructs could then be employed as probes of PIP_3_ dynamics in pharmacological inhibition experiments. With these capabilities, we provide new evidence that IQGAP1 coordinates PIP_3_ synthesis by PI3K with GTPase-dependent regulation of actin. The implications of this behavior on PI3K/PIP_3_ signaling and broader utility of this approach are discussed.

## MATERIALS AND METHODS

### Plasmids

All plasmids used for transfection were purchased from Addgene and used directly or with the indicated modifications made by Gibson Isothermal Assembly (27). Plasmids were purified according to manufacturer instructions by either Midiprep (Qiagen) or PureYield Plasmid Miniprep with Endotoxin removal wash (Promega).

### Cell culture and transfection

MCF10A human mammary gland cells were purchased from ATCC (CRL-10317). Cell line authenticity was validated by STR profiling. Cells were cultured in Mammary Epithelial Cell Growth Medium (MEGM), without GA-1000 (Lonza), but supplemented with 100 ng/mL cholera toxin (Sigma) and antibiotic (penicillin, streptomycin, and ampicillin; Gibco). Cells were transfected using a Nucleofector IIb machine (Lonza), program T-024 and Amaxa Nucleofector Solution T (Lonza). Briefly, 0.5-2 million adherent cells were treated with 1-2 mL of Trypsin-EDTA solution for 10-25 minutes until cells were detached, then cells were washed with 2-4 mL growth media supplemented with 10% fetal bovine serum to quench trypsin activity, centrifuged for 5 minutes, resuspended with 8 mL phosphate-buffered saline, centrifuged again for 5 minutes, and then resuspended in 300 *μ*L Solution T. For a single transfection, 100 *μ*L of this cell solution was incubated for 1 minute with 3 *μ*g total of the indicated plasmids prior to nucleofection. Cells were then plated onto a well of a 6-well polystyrene tissue culture dish containing 3 mL of pre-incubated MEGM. After a 24-hour recovery period, the transfected cells were then either transferred to no. 1.5 coverslips pre-coated with 1% Growth Factor Reduced Matrigel (Corning), or frozen in MEGM supplemented in 10% DMSO for later use. Cells were imaged 18-36 hours after plating on the glass slides.

### Live-cell microscopy

Imaging experiments were performed using a humidified live-cell imaging chamber (Bioptechs) warmed to 37°C in 5% CO_2_ (Okolab). Epifluorescence images were captured using an automated Nikon Ti-E inverted microscope outfitted with a 1.4 and 1.45 NA 60x objective, a motorized stage, filter wheel, perfect focus system, and a LucaR EMCCD camera (Andor) set to 2×2 binning to mitigate photobleaching in time lapse movies. Movies were recorded at 60 second framerate. High-resolution, structured illumination microscopy (SIM) images were obtained by imaging cells on Matrigel-coated glass coverslips using an OMX 3D SIM instrument.

### Intensity analysis

Live-cell images of IQGAP1-positive endosomes were segmented using a split-Bregman procedure implemented in the Squassh segmentation plugin for ImageJ (28, 29). Marker contrasts at individual compartments were calculated using: 

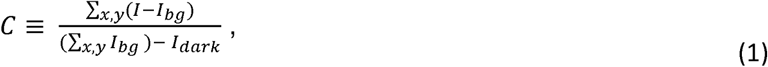

where marker-specific terms for intensity (*I*) and cytoplasmic background intensity (I_bg_) are summed over pixels (*x,y*) in the compartment-specific mask, and the fixed camera bias (the intensity in gray scale units reported by the camera in the absence of illumination, *I*_*dark*_) is subtracted. Background I_bg_ values were calculated by smoothing images with a 14 pixel rolling disk filter. *I*_*dark*_ was determined by measuring the average gray scale unit (g.s.u.) per pixel for ‘dark’ images without a sample. Similar values were found by constructing a linear fit between g.s.u. values and exposure-time and defining *I*_*dark*_ as the y-intercept of that fit. Differences in mutant localization were quantified by calculating a relative intensity contrast metric (*C*_*rel*_ ≡ *C*_*mut*_ */ C*_*wt*_), where *C*_*mut*_ and *C*_*wt*_ are the contrast of the mutant and wild-type IQGAP1 constructs, to account for potential variability in IQGAP1 contrast across transfections. C_rel_ values are reported as mean ± standard deviation. Control experiments were performed where cells were co-transfected with mVenus-IQGAP1 and FusionRed-wt-IQGAP1 in equal plasmid concentrations. C_rel_ values were close to 1 in these experiments (C_rel,mVenus-IQGAP1_ = 1.061 ± 0.129), indicating that differences in C_rel_ should correspond to differences in localization caused by domain-level mutations to the scaffold, not the choice of fluorescent protein or associated image acquisition settings.

Correlation coefficients were computed on compartments that persisted for at least 30 minutes as determined by the Squassh segmentation program (28, 29). To compute the time series of marker fluctuations, intensity trajectories of individual compartments were first normalized by *Î = I/Imax*, and then the 1^st^ derivative was computed by a 9-point first-order Savitzky-Golay filter. Dynamic cross-correlation coefficients (*r*) were then computed from this time-derivative using: 

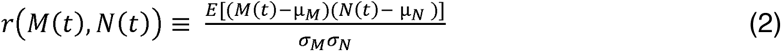

where M(t) and N(t) are time dependent marker intensity fluctuations (where fluctuations are simply the first derivative of intensity with respect to time, *μ*_M_ is the trajectory mean for marker M, and σ_M_ is the trajectory standard deviation of M. E indicates averaging across all time points in a trajectory.

Multiple regression were performed similarly using intensity time-derivative data in the R programming environment. Values reported as mean ± standard deviation.

### Multiple regression modeling

Domain-level contributions were evaluated through a series of standard multiple regression analyses. Our general procedure is illustrated in Table 1. Briefly, intensity trajectories from multiple compartments were combined to generate scatter plots of actin or Exo70 intensity as a function of mutant and wt-IQGAP1 intensity. Trajectories were down-sampled to reduce serial correlation. Regression analyses were then performed to determine if this data could be best approximated by simple linear regression model (line of best fit for a single predictor variable) or a multiple linear regression model (plane of best fit for two predictor variables). The predictive value of a regression model was quantified by calculating metrics commonly used in statistical modeling: the adjusted coefficient of determination (Adj. R^2^), the F statistic, and the change in Bayesian Information Criterion (ΔBIC). The BIC for a model is defined as

**Table 1.**

| Triple plasmid transfection | Model ($y \sim x$) | ΔBIC | Adjusted $R^2$ | F Statistic |
| --- | --- | --- | --- | --- |
| Exo70 + FusionRed-IQGAP1 + YFP-IQGAP1 | **Exo70 ~ FR IQGAP1** | **-26.6** | **0.116** | **1.69E-08** |
|  | Exo70 ~ YFP IQGAP1 | -23.6 | 0.105 | 7.65E-08 |
|  | Exo70 ~ FR IQGAP1 + YFP IQGAP1 | -22.3 | 0.117 | 7.01E-08 |
| Actin + ΔCHD + WT IQGAP1 | Actin ~ IQGAP1 | -15.4 | 0.0291 | 2.85E-06 |
| | Actin ~ ΔCHD | -82.9 | 0.117 | <2.2E-16 |
|  | **Actin ~ IQGAP1 + ΔCHD** | **-107** | **0.153** | **<2.2E-16** |
| Exo70 + ΔCHD + WT IQGAP1 | **Exo70 ~ IQGAP1** | **-43.0** | **0.0459** | **1.71E-12** |
| | Exo70 ~ ΔCHD | 6.96 | 0.0123 | 1.95E-04 |
| | Exo70 ~ IQGAP1 + ΔCHD | -45.14 | 0.0534 | 1.66E-13 |
| Actin + T1050AX2 + WT IQGAP1 | Actin ~ IQGAP1 | -2.15 | 0.0112 | 3.27E-03 |
|  | Actin ~ T1050AX2 | -57.9 | 0.0887 | <1.12E-15 |
|  | **Actin ~ IQGAP1 + T1050AX2** | **-109** | **0.161** | **<2.2E-16** |
| Exo70 + T1050AX2 + WT IQGAP1 | Exo70 ~ IQGAP1 | -66.1 | 0.106 | <2.2E-16 |
|  | **Exo70 ~ T1050AX2** | **-324** | **0.405** | **<2.2E-16** |
|  | Exo70 ~ IQGAP1 + T1050AX2 | -327 | 0.4123 | <2.2E-16 |

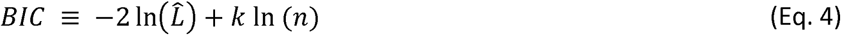

Where 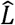 is the maximized likelihood of the model parameters, *n* is the number of degrees of freedom, and *k* is the number of fit parameters (30). The maximum likelihood function is given by -ln 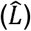 ⍰ n ln(SSR), where *SSR* is the ‘Sum of Squared Residuals’ of the regression fit. ΔBIC is calculated by subtracting the model BIC from a ‘reference BIC’ that is the BIC of a trivial model in which actin or Exo70 fluctuations are fit to a constant (e.g., actin ∼ 1). An optimal model will generally possess a minimal SSR and minimized Δ BIC. Importantly, the ΔBIC calculation includes a penalty for the degrees of freedom in a model, and therefore favors models with fewer variables. We finally note that, since the BIC is based on the maximized likelihood function, comparing ΔBIC is only appropriate for models that have been fit to the same data. The design and results of control experiments for this formalism are described in the supporting information.

## RESULTS AND DISCUSSION

The overall goal of this study is to determine how multi-functional scaffolds like IQGAP1 couple dynamic cytoskeletal and lipid membrane signaling events by coordinating the local activities of actin regulatory proteins and lipid messenger molecules, in this case Rho GTPases and the lipid messenger phosphatidylinositol (3,4,5)-trisphosphate (PIP3). We are particularly interested in how this coordination can be resolved via live-cell imaging of dynamic protein localization behaviors. Experiments employed IQGAP1-positive sorting endosomes as an experimental model. The micron size, basal confinement, low mobility, relatively short lifetime (∼1 hr) of these compartments make them highly amenable to temporal imaging analyses.

Again, we have previously shown that IQGAP1 and actin intensities correlated negatively at high-frequencies due to the temporal relationship between transient scaffold disassociation and growth of the actin networks that surround and confine endosomes to the basal cortex of polarized epithelial MCF10A cells. To further characterize this apparent negative actin regulation and probe potential coupling to other membrane development and signaling processes, we first performed a series of live-cell imaging experiments to identify and characterize the temporal hierarchy of actin regulatory proteins and PI lipid species that function within the compartments. We then examined the dynamics of IQGAP1 mutants in order to delineate how the different scaffold domains contribute to IQGAP1 intensity correlations. These assays then provided a framework to construct multiplexed statistical modeling of actin intensities and evaluate the functional interdependencies between local actin regulation and PI3K / PIP3 signaling by IQGAP1.

### IQGAP1 is coupled to Arp2/3 mediated actin polymerization and PI3K / PIP_3_ signaling

Live-cell imaging of triply-cotransfected MCF10A cells was first performed to examine dynamic relationships between fluorescent reporting constructs for IQGAP1, actin and actin regulatory proteins that have been reported to interact with IQGAP1 scaffolds (**Fig 1, Supporting Information, Fig. S1**). Among these constructs, the Arp3 component of the actin-related protein-2/3 (ARP2/3) complex displayed the most prominent dynamic relationships with actin and IQGAP1. Intensity-time trajectories of Arp3 and actin tended to follow one another (**Fig 1A, 1B**), yielding significant positive, dynamic Arp3 / actin correlation coefficients (**Fig 1C**). Arp3 also increased rapidly during actin polymerization bursts that often accompany the disassembly of the compartments and dispersal of their vesicle contents. As in our earlier studies, IQGAP1 tended to disassociate prior to these actin bursting / disassembly events. Yet, scaffold intensities correlated more negatively with Arp3 than with actin, indicating the scaffold negatively regulates Arp2/3-mediated actin polymerization. Similar behavior was also notably seen with the sortin nexin 9 (SNX9), a membrane remodeling protein that has been shown to trigger Arp2/3-dependent actin polymerization (31). The significance of this observation is discussed below.

**Figure 1.**
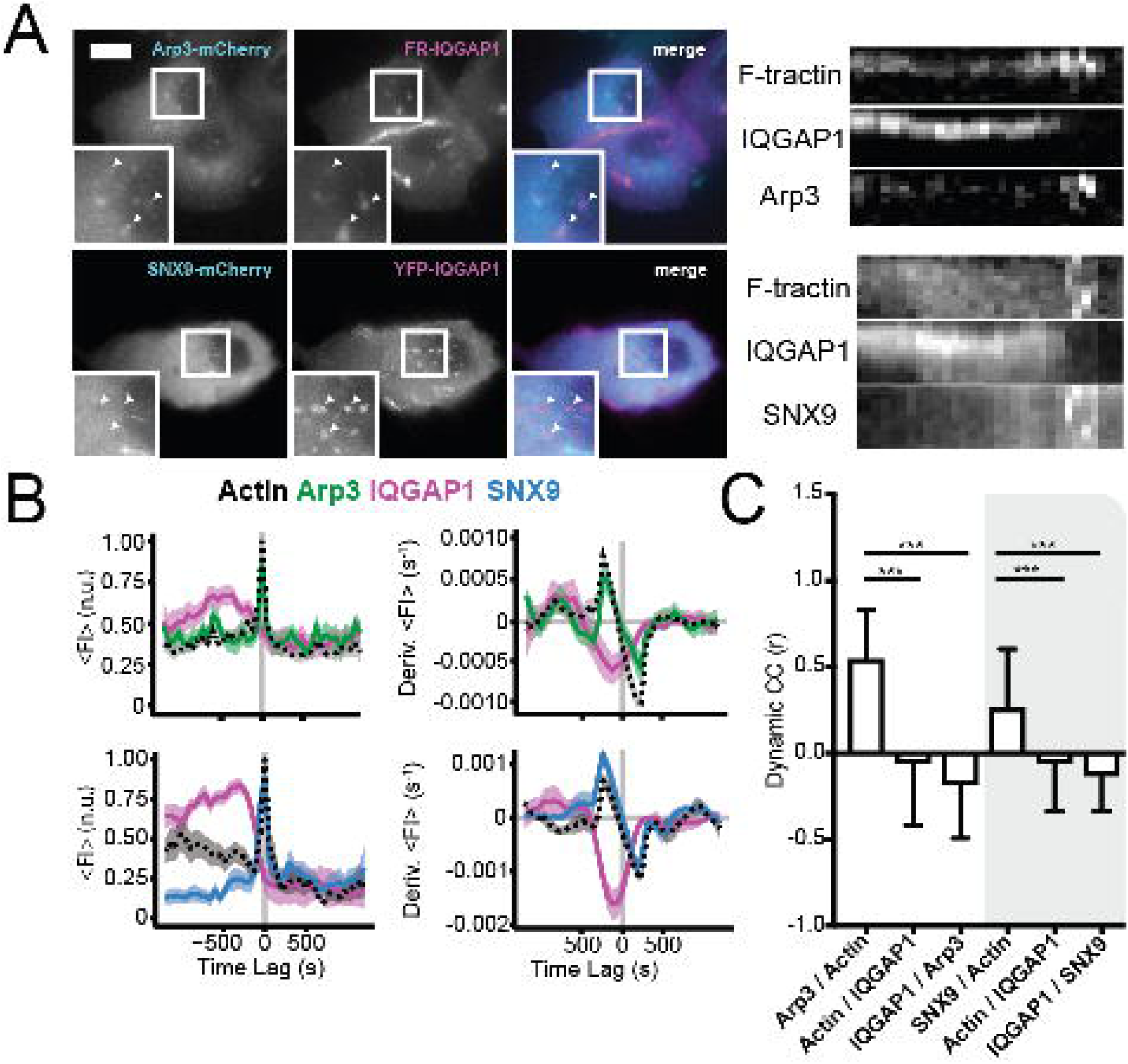
IQGAP1 disassociation prompts Arp2/3 and SNX9 dependent polymerization of actin around endosomes. (A) Live cell images and kymographs of MCF10A cells transfected with the indicated fluorescent constructs. Data are representative of n > 30 cells and three separate transfections; arrows indicate IQGAP1-containing compartments; scale bar = 10 μm. Kymographs (right panels) show rapid bursts in Arp3 and SNX9 intensity immediately after IQGAP1 dissociates from the compartments. (B) Time-dependent fluorescence intensity trajectories and their time derivatives display similar dissociation and bursting behaviors found in kymographs. Shaded area indicates standard error of the mean (for Arp3: n = 45 trajectories from 7 cells; SNX9: n = 8 trajectories from 4 cells). (C) Dynamic cross-correlation coefficients (*r*) calculated using equation 2 showing IQGAP1 is correlated negatively with Arp3 and SNX9 (*** denotes p<0.0001).

We next examined the localization behaviors of the PI species phosphatidylinositol (4,5)-bisphosphate (PIP_2_) and phosphatidylinositol (3,4,5)-trisphosphate (PIP_3_) using probes based on fluorescent protein fusions to the plextrin homology (PH) domains of PLCδ and Btk respectively. While both PI probes colocalized with IQGAP1 on the endosomes, we find the PH^_ _ _ δ^ (PIP_2_) construct to be a much more reliable probe and to exhibit more uniform labeling compared to the PH^Btk^ (PIP_3_) construct (**Fig 2 and Fig 3**). The prominent localization of PH^_ _ _ δ^ also facilitated visualization of endosome membrane morphology via live-cell structured illumination microscopy (SIM). Here, the enhanced spatial resolution of SIM revealed a range of compartment morphologies as well as the presence of membrane tubule structures that extend from the central region of the sorting compartments (**Fig 2C**). These observations are consistent with the dynamic localization of SNX9 shown in Fig 1. SNX9 is recognized for its roles in membrane remodeling and tubule formation during various stages of endosome development (31). This function presumably stems from the ability of SNX9 to sense changes in membrane curvature and promote N-Wasp-Arp2/3-dependent polymerization of branched actin networks. Moreover, the effect of SNX9 on actin polymerization has been reported to be sensitive to the membrane concentration of PIP_2_. Such behavior suggests SNX9 is responsive to both the mechanical properties and the lipid composition of endosomes, thus providing a mechanism to couple membrane development to local actin regulation. The present analyses provide new evidence that negative regulation by IQGAP1 scaffolds may be critical to this coupling and the resultant control over endosome processing.

**Figure 2.**
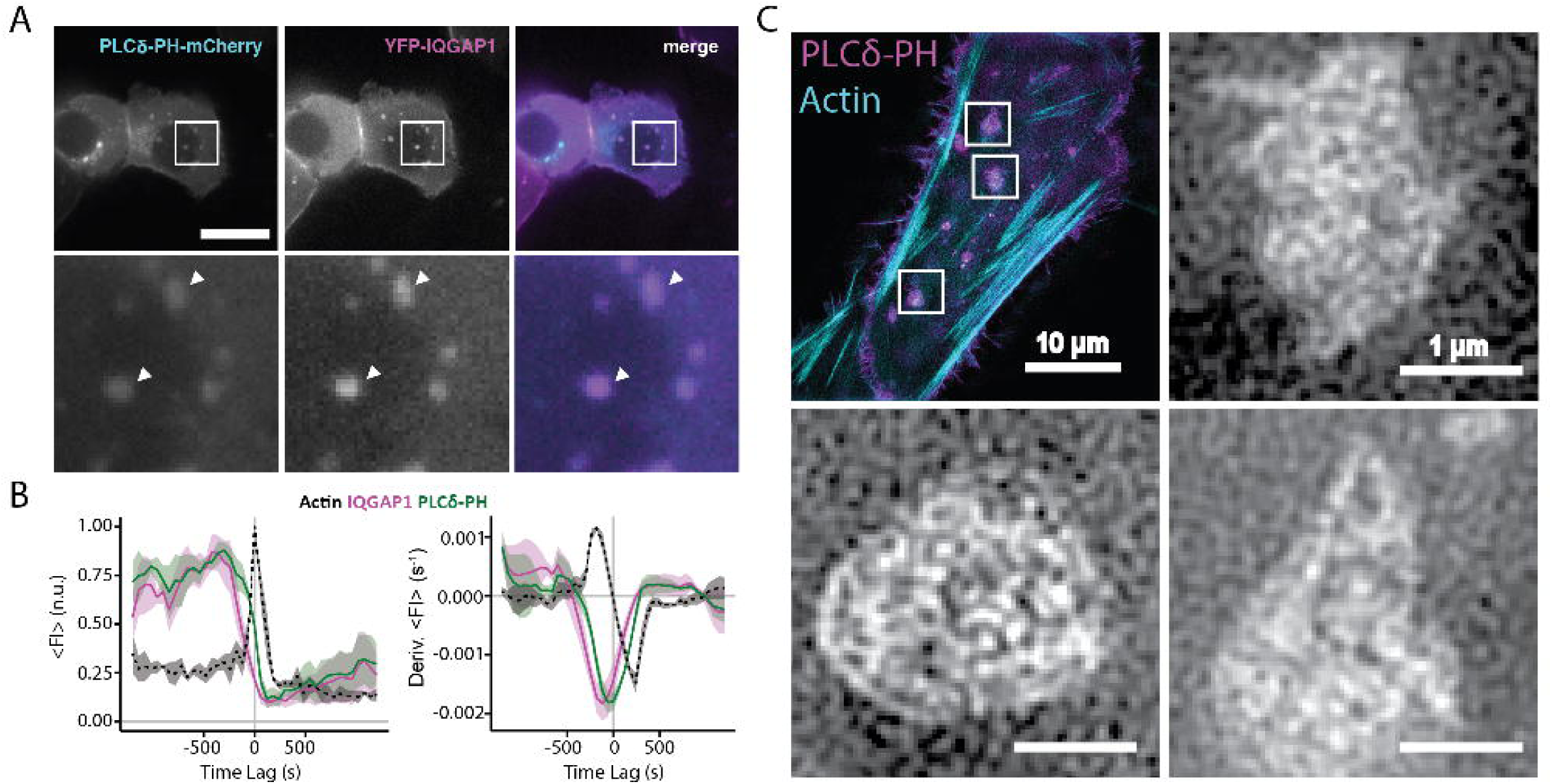
Endosomes exhibit prominent PIP_2_ localization and membrane tubule formation. (A) Colocalization of IQGAP1 with PIP_2_ lipids detected using the PH^PLC^ sensor with regards to IQGAP1 fluorescence. Scale bar = 10 μm. (B) Average fluorescence intensity and derivative of fluorescence intensity plots showing the preemptive loss of IQGAP1 and retention of PIP_2_ until the endosomes are disassembled by an actin burst (n > 30 compartments in 8 cells). (C) Membrane tubule formation on PIP_2_ - positive endosomes revealed by 3D structured illumination microscopy.

**Figure 3.**
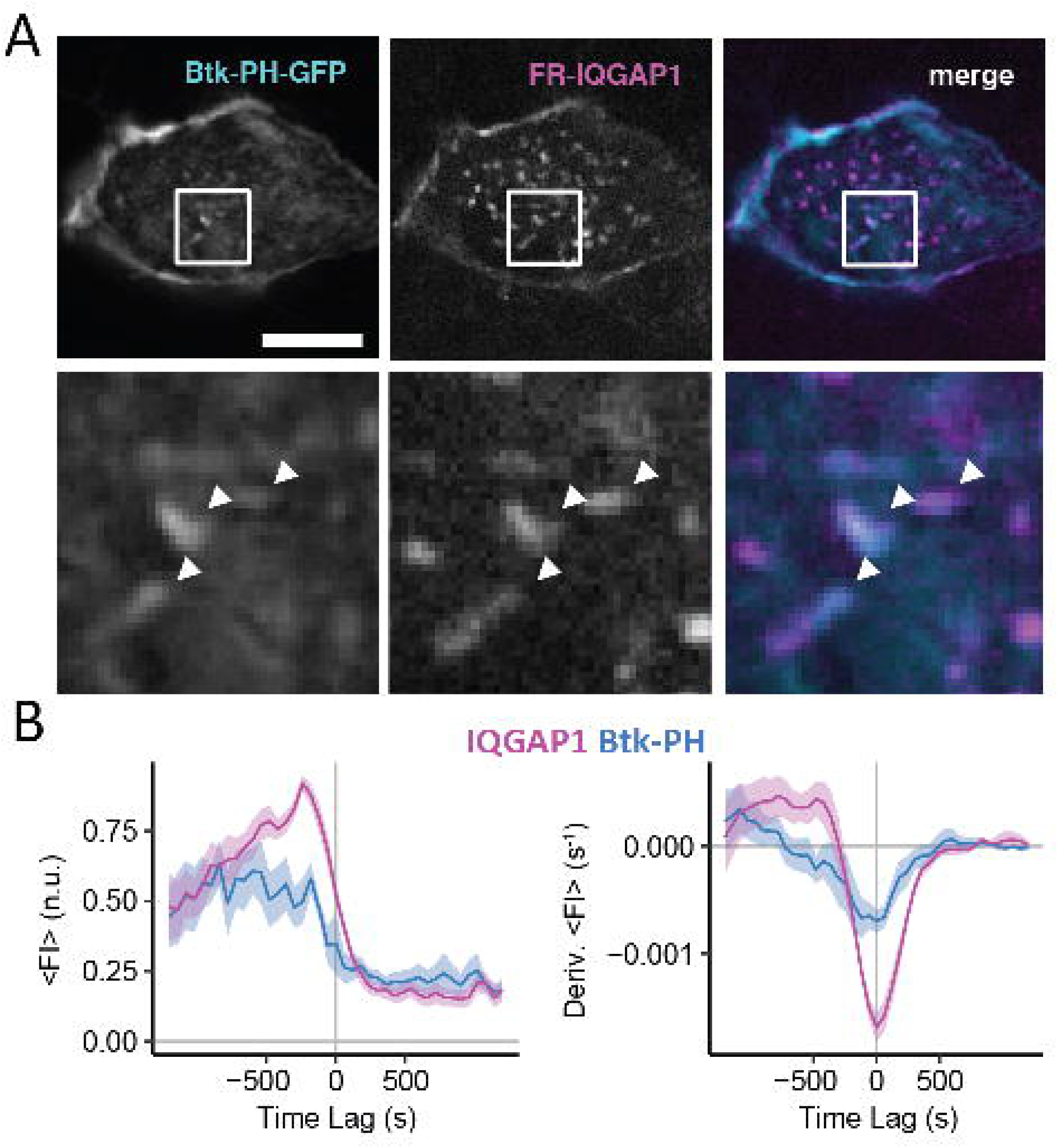
PIP_3_ exhibits weak and non-uniform localization on endosomes. (A) Example summed intensity images of IQGAP1 and PH^Btk^ colocalization. Scale bar = 10 μm. (B) Average intensity trajectories showing PH^Btk^ disassociates prior to IQGAP1. (n = 30 compartment, 8 cells).

The PH^Btk^ probe tended to express poorly in MCF10A cells, making it more difficult to find cells expressing all three probe constructs. We believe this potentially stems from inhibition of PIP_3_ signaling and its resultant cytotoxicity in MCF10A cells. As with many other PI species, PIP_3_ is generally produced transiently and at lower levels than PIP_2_. Overexpression of affinity probes like the PH^Btk^ construct may therefore be more likely to introduce inhibitory expression perturbations. Nevertheless, we were able to find examples that expressed IQGAP1, PH^Btk^ and a probe for actin (FusionRed-F-Tractin) (**Fig 3**). Endosome compartments in these cells exhibited non-uniform PH^Btk^ staining (**Fig 3A**). Analyses of average intensity-time trajectories indicate PH^Btk^ fluorescence intensity decreases prior to IQGAP1 dissociation and actin bursting, and could mean the scaffold is potentially either regulating or responding to changes in PIP_3_ levels.

Given the apparent limitations of the PH^Btk^ construct, we further probed the role of PIP_3_ using pharmacological inhibition of PI3K via the drug Wortmannin as way to inhibit local PIP_3_ synthesis. In this case, cells were triply transfected with probes for IQGAP1, actin, and PIP_2_(using PH^_ _ _ δ^). Addition of Wortmannin prompted disassociation of IQGAP1 simultaneously from multiple compartments (*t*_*1/2*_ = 12 min), followed by a gradual loss of actin (*t*_*1/2*_ = 23 min), and even more gradual dispersal and loss of PH^_ _ _ δ^/ PIP_2_ signals (*t*_*1/2*_ = 37 min). This behavior strongly implies that IQGAP1 localization and its role in actin dynamics is coupled to PI3K / PIP_3_ signaling.

### IQGAP1 exhibits multiple tethering modes

We next examined how different scaffold domains contribute to IQGAP1 localization by comparing the intensities and contrast of fluorescent protein-tagged scaffolds to IQGAP1 domain deletion and point mutation constructs. Mutant localization was evaluated by quantifying potential losses in contrast relative to a wild type scaffold that incorporated a N-terminal fluorescent protein fusion (C_rel_) (*Method Section* and the *Supporting Information, Fig S2*). Key mutant scaffolds included a domain deletion mutant (ΔCHD), where the first 175 N-terminal amino acids of IQGAP1 corresponding to the actin-binding CHD domain were removed, and a previously-investigated T1050AX2 mutant, in which a substitution, TVI (1050-1052) -> AAAKMVVSFNEIVTGNPAAA, has been introduced in the GRD (32, 33). The T1050AX2 mutant is thought to represent a constitutively active form of IQGAP1 that behaves as if IQGAP1 were constitutively bound to active (GTP-bound) Cdc42, although, counterintuitively, this mutation eliminates IQGAP1 co-immunoprecipitation with Cdc42 and Rac1 GTPases. We also examined a scaffold incorporating a C-terminal fluorescent protein tag and a GRD domain mutant (GRD6A) (34), where six alanine substitutions (N1011A, N1015A, Q1016A, R1017A, E1018A, and T1291A) were made in the GRD of the full-length scaffold that have been shown to disrupt interactions with Rho GTPases.

Comparisons of mutant constructs suggest that the C-terminal GRD–RGCT scaffold domains are the key drivers of IQGAP1 localization to the endosomes (Supporting Information, Fig S2). This dominant contribution is reflected by the nearly ten-fold drop in contrast for wild type constructs incorporating a C-terminal fluorescent protein fusion (C_rel, wt-IQGAP-FP_ = 0.176 ± 0.0418) and an even larger decrease for the GRD6A mutant (C_rel, GRD6A_ = 0.108 ± 0.0478), which should exhibit impaired interactions with Rho GTPases. Of note, the relative contrast of the presumed ‘constitutively active’ T1050AX2 mutant was quite high compared to the other mutants (C_rel_ = 0.632 ± 0.170), indicating that, despite the potential loss of GTPase associations that has been reported for this construct, the T1050AX2 mutation does not impair scaffold localization appreciably. The T1050AX2 was also found to localize to some IQGAP1 negative endosomes compartments. The dynamic behaviors underlying this localization heterogeneity are discussed below.

Despite the predominance of the C-terminal GRD-RGCT domains, the actin-binding CHD at the N-terminus of the scaffold still appears to play an appreciable role in IQGAP1 localization. In particular, the contrast of the ΔCHD contrast is roughly half of that of the FP-IQGAP1 construct (C_rel, ΔCHD =_ 0.507 ± 0.196), indicating that CHD / actin interactions confer substantial affinity to the endosomes. Moreover, a double mutant construct (ΔCHD-GRD6A) where the CHD was deleted from the GRD6A mutant did not localize to the compartments either (C_rel, ΔCHD-GRD6A_ = 0.0478 ± 0.0972) (Supporting Information, Fig. S3). In this case, compartment-localized voids are found in summed intensity images, suggesting that the modest localization of the GRD6A single mutant can be attributed to the CHD. A double mutant (ΔCHD-T1050AX2) also exhibited dramatically impaired contrast (C_rel_ = 0.145 ± 0.104)(Fig. S1). Together, these analyses indicate that, while IQGAP1 binding to the endosome compartments is largely governed by the C-terminal GRD-RGCT domains, localization is assisted greatly by the actin binding CHD. Retention of the ΔCHD-T1050AX2 double mutant is suggestive of a third interaction, likely between the scaffold and the membrane. Regardless, these behaviors suggest that time-dependent scaffold localization intensities will ultimately be governed by the dynamics of at least two scaffold domains and their associated actin or endosome tethering interactions.

### The GTPase activity regulates scaffold tethering at temporal high frequencies

Time-lapse live-cell imaging was next used to examine differences in the dynamic properties of the wild type scaffold and the ΔCHD and T1050AX2 mutants (Fig 5). Compartment dynamics was monitored in 3-color experiments using MCF10A cells expressing FusionRed-wt-IQGAP1 (*hereafter IQGAP1*), a mutant mVenus-tagged IQGAP1 (*either ΔCHD, or T1050AX2*), and a mTurquoise2-tagged label for either actin (*F-Tractin*) or a membrane protein (*Exo70*) that we have found to exhibit high image contrast in the compartment. Marker intensities were imaged at 1 frame per minute for 90 min. For experiments incorporating F-Tractin, the relative timing of mutant and IQGAP1 binding and unbinding was evaluated by aligning and averaging intensity-time trajectories from multiple compartments using the peak in actin intensity that occurs during compartment disassembly / actin bursting events as a registration point (Fig 5A, 5B). Since Exo70 signals disappear during compartment disassembly, analyses of membrane-associated dynamics were registered using the minimum of the first derivative of Exo70 signals (Fig 5D, 5E).

**Figure 4.**
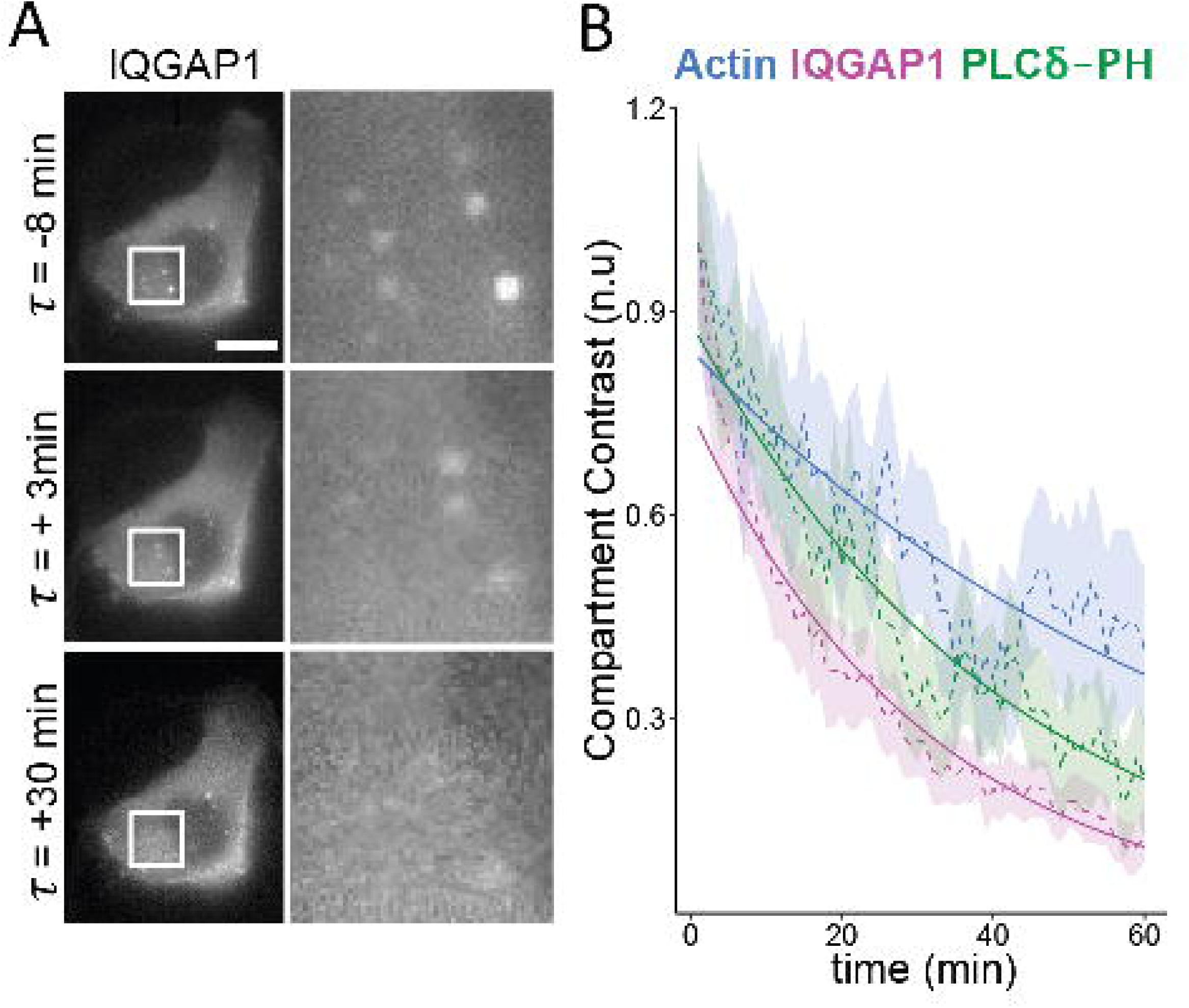
PI3K inhibition prompts IQGAP1 disassociation and subsequent disassembly of the compartments. (A) Epifluorescence images monitoring the disassociation of IQGAP1 from endosomes in Wortmannin treated MCF10A cells. Scale bar = 10 μm. (B) Intensity-time plots showing the loss of IQGAP1, PIP_2_, and actin signals in Wortmannin treated cells. Dashed lines, shaded area indicates SEM. Solid lines indicate exponential fits. (n=45 compartments, 7 cells).

**Figure 5.**
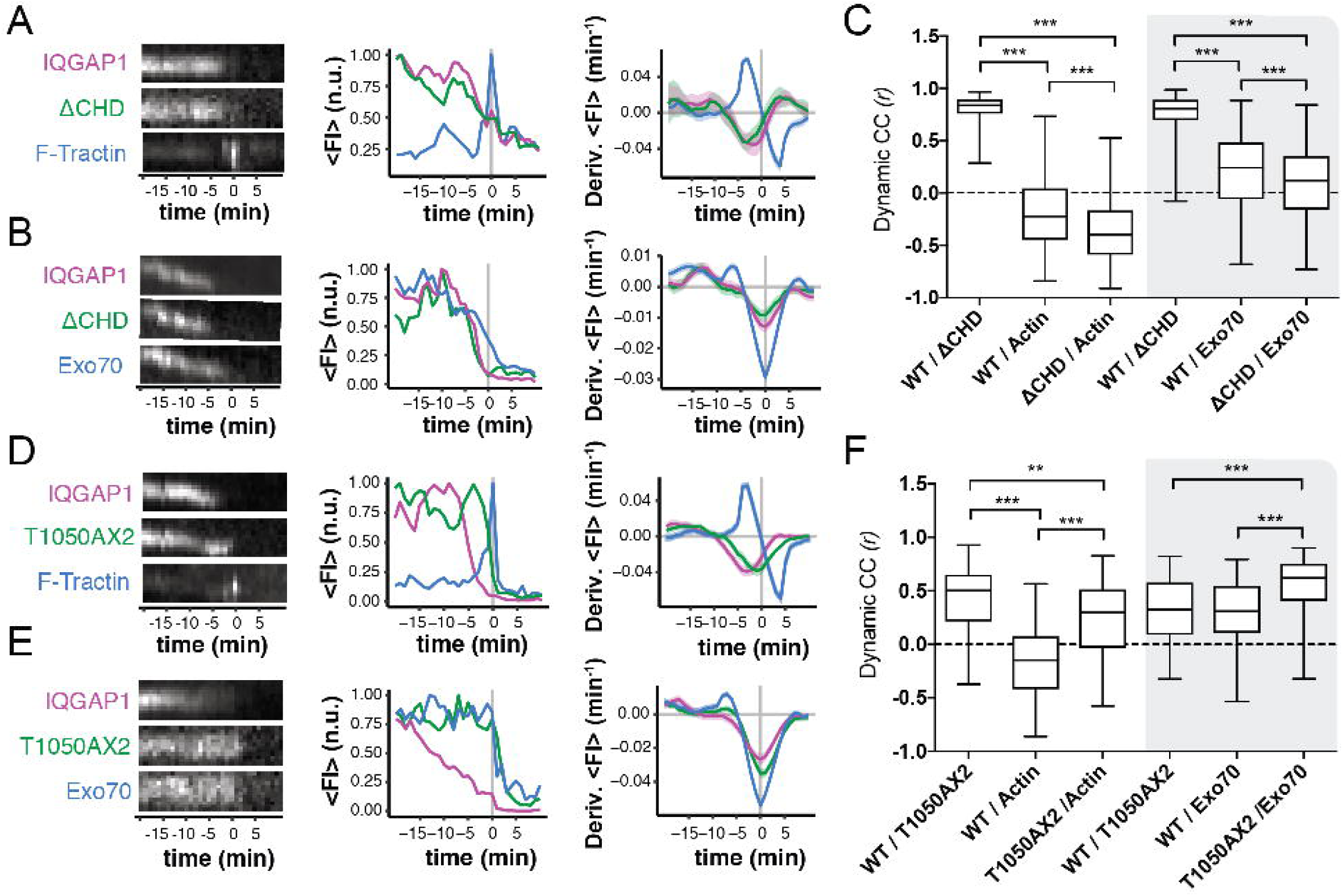
(A) (*left*) Kymographs and their corresponding normalized intensity-time trajectories (*middle*) for the ΔCHD mutant, wild type IQGAP1, and F-Tractin, intensity trajectory for a representative compartment with an actin burst, and (*right*) time-derivative plots averaged over multiple trajectories showing mean and SEM for each marker (n = 14 compartments). (B) Intensity time data for ΔCHD, WT IQGAP1, and Exo70 aligned using the minimum of the derivative of the Exo70 signal. (n = 241 inflection points). (C) Dynamic cross correlation coefficients (*r*) calculated for combination of wild type and mutant IQGAP1 scaffolds, and either actin or Exo70 (Actin: n = 234 compartments, 19 cells; Exo70: 183 compartments, 17 cells; *** denotes p<0.0001). (D-F) Intensity time and cross correlation plots for the T1050AX2 mutant. (Actin: n = 71 bursts, 354 compartments, 11 cells; Exo70: n = 163 inflection points, 363 compartments, and 12 cells; ** denotes p=0.005).

Consistent with the CHD playing a secondary role in scaffold localization, time-dependent fluorescent intensity trajectories of the ΔCHD mutant were quite similar to those of wild type IQGAP1 construct (Fig. 5A, 5B). The ΔCHD mutant also disassociated from the compartment prior to actin bursting events similarly to IQGAP1. This behavior yields negative derivatives in the fluorescence intensity (FI) with respect to time, as in Figure 1. ΔCHD and IQGAP1 were also found to dissociate prior to the loss of Exo70 signal (Fig. 5B). Overall, only minor deviations from wild type behaviors were observed. In contrast, fluorescence T1050AX2 mutant did not dissociate prior to actin bursting events or loss of Exo70 (Figure 5D-E), but was, instead, retained until compartments were fully disassembled by an actin burst. This behavior implies that T1050AX2 mutant is anchored tightly to membrane components of the endosome compartment. The mutation in this ‘constitutively active’ construct therefore likely disrupts the ability of factors to modulate scaffold-membrane tethering. The inability of the T1050AX2 to bind Cdc42 or Rac1 implicates Rho GTPases in this regulation. Of note, the retention of the T1050AX2 mutant during compartment disassembly also explains the heterogeneity in wild type / mutant colocalization (Supporting Information Fig. S2), since this behavior will cause mutant signals to persist after IQGAP1 dissociation.

Analyses of dynamic intensity cross correlations also revealed important differences between the binding modes of the ΔCHD and T1050AX2 mutants (Fig 5C, 5F). In addition to highlighting a role for the CHD, several of these differences further implicate Rho GTPases in the localization of wt-IQGAP1 scaffolds. The time dependent intensities of the ΔCHD and IQGAP1 were highly correlated (*r*_Δ*CHD, IQGAP1*_ = 0.810 ± 0.120). While this behavior further suggests the CHD plays a minority role in IQGAP1 localization, as with the contrast measurements, this domain still appears to have an appreciable effect on IQGAP1 localization dynamics. In particular, ΔCHD intensities are more negatively correlated with actin intensity compared to IQGAP1 (*r*_Δ*CHD, actin*_ = −0.365 ± 0.294, and *r*_*IQGAP1, actin*_ = −0.191 ± 0.326, p < 0.0001) (Fig. 5C). These correlation enhancements indicate that actin binding by the N-terminal CHD obscures abilities to resolve negative IQGAP1 / scaffold regulatory interactions that occur at relatively high frequencies (∼1 minute timescales), perhaps by preventing or slowing total IQGAP1 disassociation from the compartment. Of note, analyses of correlations during the growth and disassembly using Hidden Markov Modeling (HMM) show that this negative regulation is a persistent feature of IQGAP1-mediated actin regulation on endosomes and occurs during both the growth and disassembly phases of the compartments (Supporting Information, Fig S4).

In contrast to the ΔCHD, intensity fluctuations of the T1050AX2 mutant correlate positively instead of negatively with actin (*r*_*T1050AX2, actin*_ = 0.234 ± 0.359) (Fig. 5F). The average correlation between T1050AX2 and Exo70 intensities (*r*_*T1050AX2, Exo70*_ = 0.563 ± 0.248) was also significantly higher than between IQGAP1 and Exo70 (*r*_*IQGAP1, Exo70*_ = 0.277 ± 0.319). Considering again the lack of Rho GTPase binding by the T1050AX2 mutant, these results suggest that the negative correlations between IQGAP1 binding and actin polymerization, and the actin inhibitory mechanism we attribute to this behavior, depend on compartment-associated GTPases and their local effectors. The strong positive correlation between Exo70 and T1050AX2 intensities suggests that scaffold localization of this mutant may also be driven by other membrane-associated compartment components. Analyses indicating this behavior stems from the combined actin and membrane tethering functions of the scaffold are discussed in the next section.

### Multiple regression modeling delineates contributions of different scaffold tethering modes

The idea that IQGAP1 localization depends on multiple domains naturally raises the possibility that this scaffold associates with the endosome compartments in different states where it is tethered via actin / CHD interactions, direct or indirect GRD - membrane interactions, or a combination of these interactions. If so, dynamic IQGAP1 intensities could represent a weighted sum of fluorescent intensities generated due to the association and disassociation of each of these subspecies. With the goal of further teasing out the identity and contribution of these interactions to scaffold localization, we next tested whether the ΔCHD and T1050AX2 mutants approximate the localization behavior of different scaffold intermediates by evaluating a series of simple single and multiple regression models that assumed actin or Exo70 signals depend on a single mutant, the wild type scaffold, or a combination of a mutant and the wild type IQGAP1 scaffold (Table 1). Regression models were evaluated by calculating reduced R^2^ values along with changes in a parameter called a Bayesian Information Criterion (ΔBIC), which evaluates model fitness relative to a base model that assumes signals are uncorrelated (*see the Method Section*). For reference, model differences yielding ΔΔBIC > 10 is generally considered to be very strong evidence for preferring one model over another, while ΔΔBIC ≤ 2 is considered negligible (30).

Calculated values for both ΔBIC and R^2^ for the simple linear model [actin ∼ IQGAP1] are quite low, indicating that IQGAP1 carries very little information regarding actin intensity on its own (Table 1). The very weak dependence of actin on IQGAP1 in regression models notably persisted in multiple experiments with different combinations of markers. We attribute this behavior to the dependence of IQGAP1 localization and regulatory activity on mixtures of different scaffold domains and interactions. A similar linear model [actin ∼ ΔCHD] that assumes actin depends on the remaining regulatory domains in the ΔCHD provides better regression fits to the actin data. Yet, the multiple regression model incorporating dependencies on both constructs, [actin *∼* ΔCHD + IQGAP1] yields the best description of actin intensities. Likewise, substantially improved fits are achieved with a multiple regression model incorporating T1050AX2 with IQGAP1: [actin *∼* T1050AX2 + IQGAP1]. In other words, the full length IQGAP1 scaffold is only a useful descriptor of actin dynamics when the model also accounts for correlations between actin and a mutant that is either missing actin binding, or GTPase regulation. Considering the different correlative behaviors of both mutants in Fig 5, we interpret these responses to signify that the ΔCHD and T1050AX2 mutants capture different key endosome-scaffold interactions / intermediate state behaviors of IQGAP1. The improved utility to the wild type scaffold in multiple regression therefore suggests actin signals are dependent on both of these interactions as a minimum (*i.e., the inclusion of one interaction in a model via a mutant allows the other*(*s*) *to be captured by the wild type scaffold*).

Interestingly, opposite trends occur with the membrane protein Exo70 (Table 1). Here, IQGAP1 captures more of the variation in Exo70 intensity than the ΔCHD (ΔBIC = −43 vs −7, respectively). Very little benefit is gained by considering a combination of ΔCHD and wt-IQGAP1 signals in a model describing Exo70 (ΔBIC = −45). Similarly, the actin and membrane binding T1050AX2 mutant captures Exo70 fluctuation quite well on its own, and little is gained by incorporating the wild type scaffold into the model. These trends indicate that actin binding by IQGAP1 is linked to temporal changes in membrane signals (reflected by Exo70). However, for the wild-type scaffold, this link to the membrane is affected strongly by GTPase signaling in the compartment.

### Actin inhibition by IQGAP1 is linked to PI3K-PIP_3_ signaling

Finally, to gain more insight into scaffold interactions at the endosome membrane, we next evaluated whether the membrane binding of IQGAP1 could be associated with specific phosphoinositide signals instead of the T1050AX2 mutant. In this case, MCF10A cells were triply transfected with the F-Tractin probe, IQGAP1, and the PH^_ _ _ δ^ probe for PIP_2_. As with the mutants, while IQGAP1 is a poor descriptor of actin dynamics on its own, the PIP_2_ marker PH^_ _ _ δ^ is able to capture actin dynamics similarly to the T1050AX2 mutant (Table 2). Yet, the combination of both markers yield the lowest ΔBIC, indicating this is best of the three models of actin dynamics. As with the mutant analyses above, the improved utility of the IQGAP1 to capture variations in endosomal actin intensity when models account for fluctuations in PIP_2_ membrane signals, indicates that, like the T1050AX2 mutant, the PH^_ _ _ δ^ can be used to ‘draw out’ the negative actin regulatory function of IQGAP1, that again we associate with GRD coupling to GTPase signaling. The PH^Btk^ construct has limited utility for these analyses due to its poor labeling and cytotoxicity, as described above. Nevertheless, the role of PI3K / PIP_3_ signaling could be probed via pharmacological inhibition with Wortmannin. In this case, even dilute concentration of the drug is shown to only affect the utility of IQGAP1 to describe actin dynamics in the multiple regression model. As a result, the multiple regression model is no better than the simple linear model [actin ∼ PLCδ]. This sensitivity to PI3K inhibition suggests the production of PIP_3_ lipid messengers is specifically couples to, and likely even potentiates, the negative IQGAP1 / actin regulation that is driven by the GRD. IQGAP1 may therefore operate as a hub between PI3K signaling and GTPase – mediated actin regulation.

**Table 2.**

| Model (y ~ x) | Wortmannin (pM) | ΔBIC | Adjusted $R^2$ | F statistic |
| --- | --- | --- | --- | --- |
| Actin ~ WT IQGAP1 | 0 | 5.42 | -0.00444 | 0.984 |
| Actin ~ PLCδ | 0 | -35.6 | 0.162 | 1.85E-08 |
| **Actin ~ WT IQGAP1 + PLCδ** | 0 | **-53.3** | **0.240** | **1.84E-14** |
| Actin ~ WT IQGAP1 | 100 | 3.94 | -0.00454 | 0.446 |
| **Actin ~ PLCδ** | 100 | **-48.3** | **0.1816** | **1.22E-05** |
| Actin ~ WT IQGAP1 + PLCδ | 100 | -45.6 | 0.18845 | 3.09E-05 |
| Actin ~ WT IQGAP1 | 500 | -5.4 | 0.0396 | 1.02E+03 |
| **Actin ~ PLCδ** | 500 | **-48.3** | **0.194** | **2.91E-13** |
| Actin ~ WT IQGAP1 + PLCδ | 500 | -45.6 | 0.200 | 7.17E-13 |

## Conclusion

The abilities of scaffolds like IQGAP1 to assemble, anchor, and regulate multiple protein complexes locally at specific subcellular membrane compartments are critical to a host of intracellular signaling and regulatory networks, particularly those requiring the recruitment and dynamic coordination of multiple messenger molecules, low affinity signaling intermediates and regulatory proteins such as PI signaling pathways. Nevertheless, resolving the functional regulatory roles of many scaffolds remains challenging given the spectrum of interactions that can occur in these complexes. This work demonstrates that dynamic correlation and statistical modeling analyses can be leveraged to quantify the temporal contributions of specific scaffold domains and compartment binding modes observed in live cell imaging experiments. While using IQGAP1 positive endosomes as a test case, these methods revealed that IQGAP1 scaffold can transition between different actin and membrane tethered states. We provide evidence that this transitioning allows IQGAP1 to coordinate the growth and disassembly of endosome-specific actin networks with local PI signaling, and potentially endosomal membrane processing (e.g., tubule formation).

Our results indicate that IQGAP1 inhibits actin polymerization in at least one of its tethered states, likely via negative regulation of SNX9 and Arp2/3. Comparisons of mutant scaffold constructs suggest direct actin binding by the N-terminal CHD of IQGAP1 is somewhat dispensable and does have a direct role in this negative regulation. However, CHD interactions with actin do appear to help retain the scaffold within the endosomal actin network, which could potentially reinforce suppressive actin interactions driven by C-terminal scaffold domains. Correlation analyses and regression modeling also indicate the GTPase-binding function of GRD region of the scaffold constitutes a dominant regulatory axis to control IQGAP1 tethering and actin polymerization. The difference in the membrane retention behaviors of the T1050AX2 mutant and wild type scaffold also suggests the GRD domain specifically modulates scaffold tethering to the endosome membrane.

While these results narrow the possibilities, the mechanism underlying IQGAP1 inhibition of transient actin polymerization however is still not fully resolved. Considering the ability of IQGAP1 to stabilize the GTP bound state of Rho GTPases (35), one possibility is that IQGAP1 localization restricts local GTPase activity by sequestering activated GTP-bound forms of Rac1 or Cdc42. Our data indicating the GRD mediates membrane tethering and observations of anti-correlated IQGAP1 / SNX9 localization raises the alternative possibility that IQGAP1 interactions with membrane lipids or membrane proteins inhibit SNX9 localization and subsequent activation of Arp2/3 mediated actin polymerization. Choi *et al* have notably reported that the C-terminal domains of IQGAP1 can associate directly with PIP_2_ (19), which is, again, a known effector of SNX9. It is therefore possible that the GRD regulates membrane tethering by modulating direct scaffold PIP_2_ interactions, potentially with other associations.

Finally, despite the difficulties monitoring PIP_3_ signaling events directly in this setting, analyses of scaffold binding modes in multiple regression models provide new evidence that actin regulating GRD-dependent binding / dissociation transitions are coupled to PI3K / PIP_3_ signaling. Again, we believe these transitions underlie the negative actin regulation behaviors of the scaffold and yield the negative actin / IQGAP1 intensity correlations. Extensive studies have shown that PIP_3_ signaling can activate downstream actin polymerization via Rho GTPase signaling. However, the present results now implicate IQGAP1 scaffolds at a key element in a regulatory hub that controls this coupling. Positive feedback from Rho GTPases to PI3K is also widely recognizes as important to PIP_3_ dependent signaling and actin polymerization. Given the present behaviors, we propose that the negative regulatory behavior of IQGAP1 and its potential ties to GTPase signaling may provide a mechanism to suppress this positive feedback in order to prevent overrun PI3K signaling at endosome and other signaling sites. IQGAP1 may also help tie PIP_3_ signaling to other local membrane processing and sculpting events, potentially via SNX9. Although further investigation is ultimately required, resolving these issues and the functions of many other scaffolds within local subcellular compartments will likely benefit appreciably from abilities to ‘demultiplex’ the integrated dynamics of scaffolds at the domain level.

## Acknowledgments

We are grateful for helpful conversations with David Sacks, David Worthylake, and James McNew, and funding support from the NIH (5R01GM106027-03) and the Welch Foundation (C-1625) to MRD.

## Conflict of Interest

The authors declare that they have no conflicts of interest with the contents of this article. The content is solely the responsibility of the authors and does not necessarily represent the official views of the National Institutes of Health.

