## Supplemental Figures for "Demultiplexing overlapping signaling scaffold functions to probe lipid messenger coupling to cytoskeletal dynamics"

**Supporting Figures S1 – S4**

**Supporting Table 1**

**Supporting Methods for Multiple Regression Control Experiments**

**Supporting Figure S1**

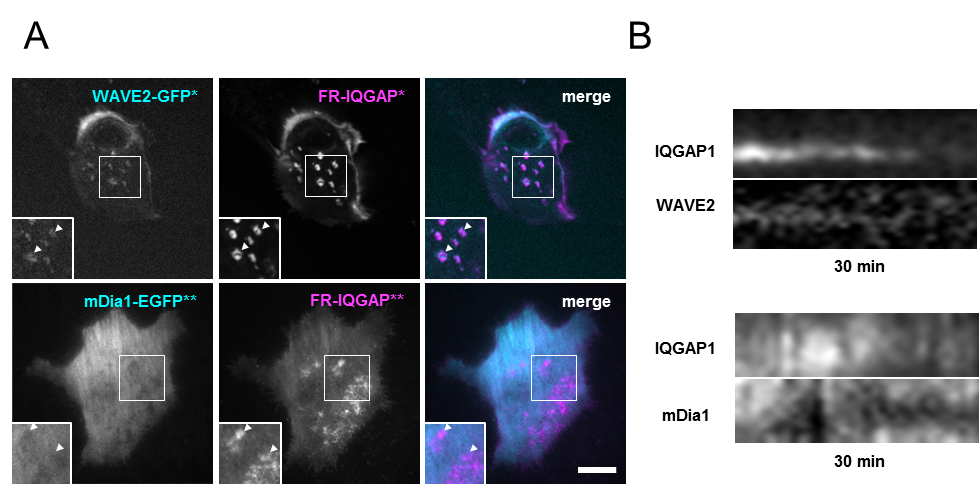

**Supporting Fig S1.** **Localization behavior of the IQGAP1 interacting proteins WAVE2 and mDia1. A)** A GFP fusion construct to the WASP family Verprolin-homologous protein 2 (WAVE2-GFP) localizes weakly to the endosome compartments while actin nucleation protein Diaphanous-related formin-1 (mDia1) does not localize at all. The WAVE2 and IQGAP1 images are summed intensity images of a cell over a 60 min movie. mDia1 images were collected using total internal reflection fluorescence microscopy (TIRFM). Scale bars are 10 µM. B) Kymographs of individual compartments during the disassociation of IQGAP1.

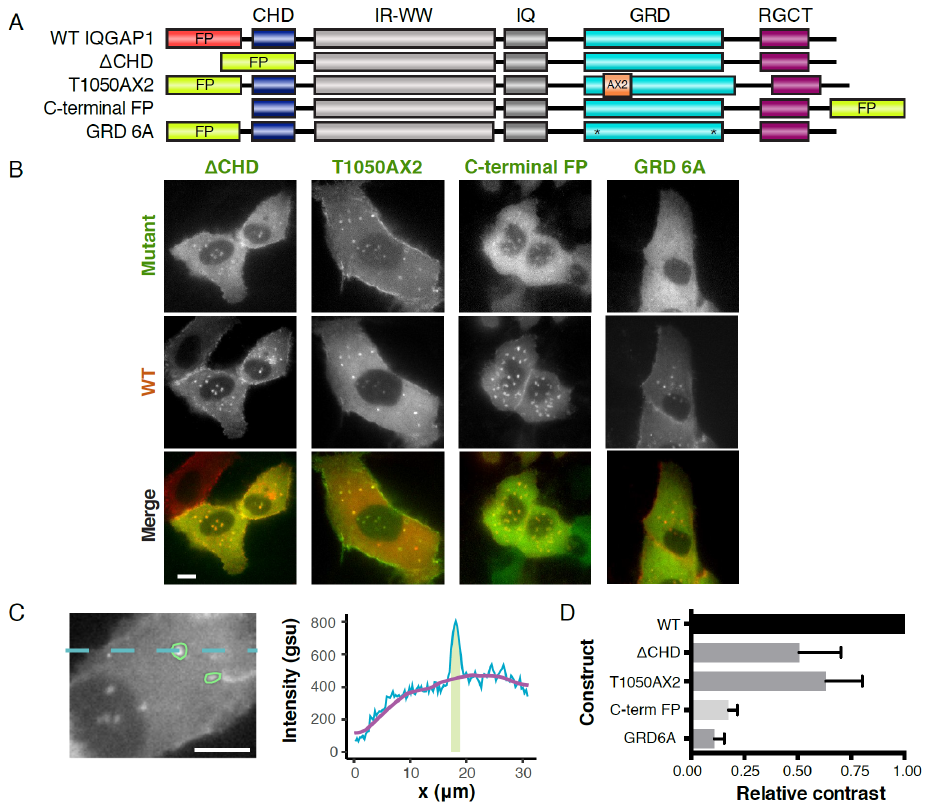
**Supporting Figure S2**

**Supporting Fig S2. Localization of IQGAP1 mutants.** MCF10As were co-transfected with mVenus-mutant and FusionRed-wt-IQGAP1, recovered for 24 hours before seeding onto Matrigel-coated coverslips. Cells were imaged 24 hours later in two fluorescent channels with an epifluorescent microscope at 60X magnification. **A)** Schematics of the domain structure for the wild type and mutant constructs drawn to scale with respect to amino acid sequence length. Red colored FP = FusionRed.  Green colored FP = mVenus.   * denotes the location of alanine substitutions in the Ex domain for the GRD6A mutant which are believed to be proximal in the 3-dimensional, folded IQGAP1 structure. **B)** Representative images of MCF10A cells co-transfected with 1 μg of the indicated mVenus-labeled mutant IQGAP1 and 1 μg wild-type scaffold (FusionRed-wt-IQGAP1).  Scale bar is 10 μm.  **C)** Schematic depiction of the contrast measurements. Compartment masks (green outlines) are generated by image segmentation on the IQGAP1 channel.  Masks are excluded if they overlap with a user-drawn nuclear mask since exclusion of the cytoplasm in this region results in circumstantially low background levels.  Average cytoplasmic background levels (purple line) were determined from an image that was smoothed using a 15 pixel rolling ball filter.  Contrast is calculated by comparing the average intensity of pixels within a mask (green) to cytoplasmic background intensities.  **D)** Relative contrast measurements for IQGAP1 constructs.  Relative contrast values were calculated from measurements of 40 to 148 distinct compartments averaged over 6 to 9 cells.

**Supporting Figure S3.**

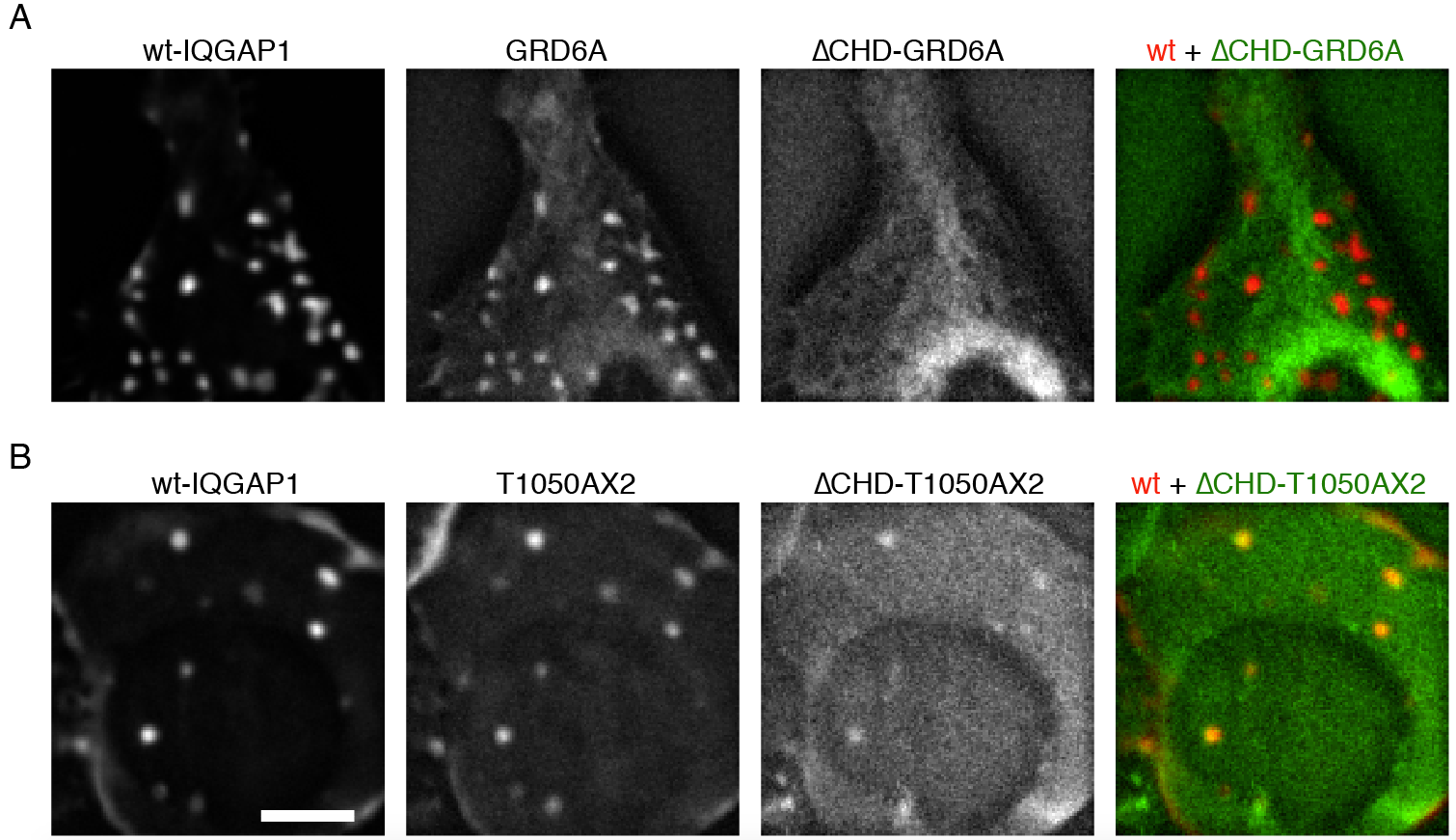

**Supporting Fig S3. Calponin Homology Domain (CHD) is necessary for GRD6A mutant localization and plays a positive contributing role in T1050AX2 mutant localization.** MCF10As were co-transfected with three IQGAP1 constructs expressing wild type, single mutant, and double mutant fused to FusionRed, mVenus, and mTFP1, respectively. Three color 60 second framerate movies were background subtracted and images were prepared with a 60 frame summed intensity projection. **A)** Single mutant GRD6A localizes poorly, whereas double mutant ∆CHD-GRD6A fails to localize, forming voids instead. Merged image (*right*) shows the double mutant is completely excluded from IQGAP1-positive compartments. **B)** Single mutant T1050AX2 localizes and ∆CHD-T1050AX2 localizes weakly to the compartments. Representative images are provided. Scale bar indicates 10 µm.

**Supporting Figure S4.**

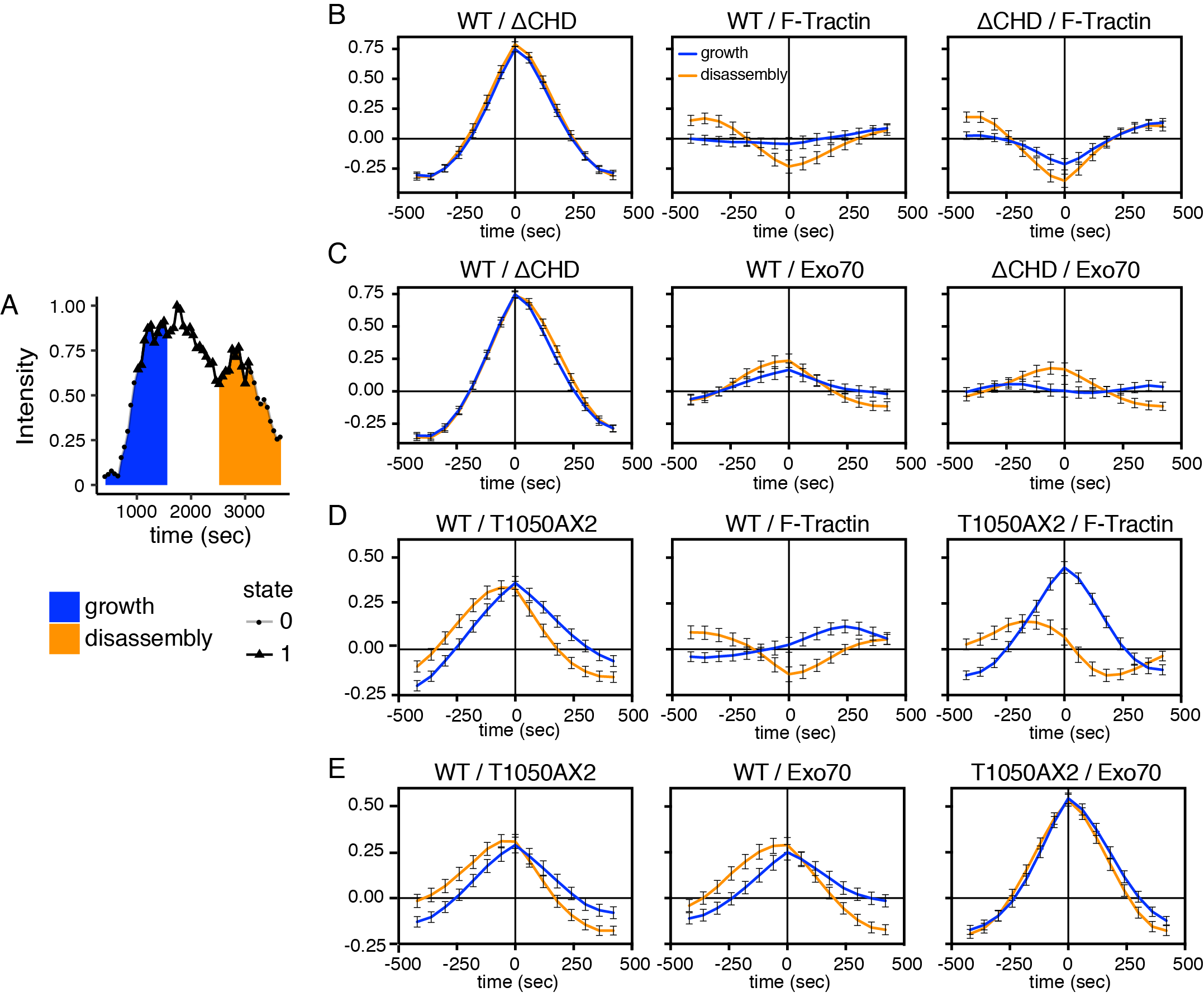

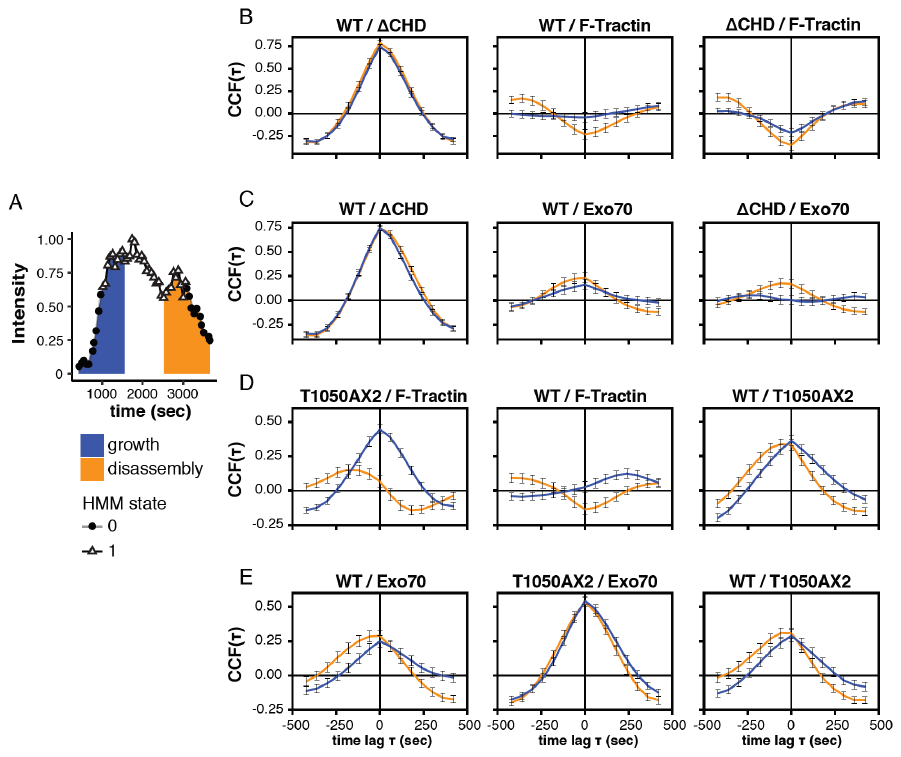

**Supporting Fig S4. Actin binding by the CHD obscures the underlying negative actin regulation by the GRD-RGCT.**  Dynamic intensity trajectories were also evaluated using Hidden Markov Modeling (HMM) as an objective method to segment compartment trajectories into segments characterized by persistent ‘growth’ or ‘disassembly’ of IQGAP1. A) Segments of compartment intensity trajectories were classified using a two-state Hidden Markov to identify how compartments transition between states characterized by ‘high’ or ‘low’ intensities of wild type IQGAP1 constructs. Normalized wt-IQGAP1 intensities were first discretized into 8 bins. The high and low intensity state distributions and transition probabilities were trained using the Baum-Welch algorithm in MATLAB, and these estimates were used by the Viterbi algorithm to classify IQGAP1 trajectories into high or low hidden states. Growth and disassembly were taken as the 10 minutes centered around low-to-high and high-to-low transitions.. **B)-E)** Cross-correlation functions, *CCF(τ) = (f⋆g)(τ),* for pairs of proteins in compartments delineated by growth and disassembly maturation states.  **B)** ΔCHD, wt-IQGAP1, and F-Tractin transfection experiments corresponding to Fig 5A.  **C)** ΔCHD, wt-IQGAP1, and Exo70 experiment corresponding to Fig 5B.  **D)** T1050AX2, wt-IQGAP1, and F-Tractin experiment corresponding to Fig 5D.  **E)** T1050AX2, wt-IQGAP1, and Exo70 experiment corresponding to Fig 5E. Error bars represent SEM.

**Supporting Table 1**
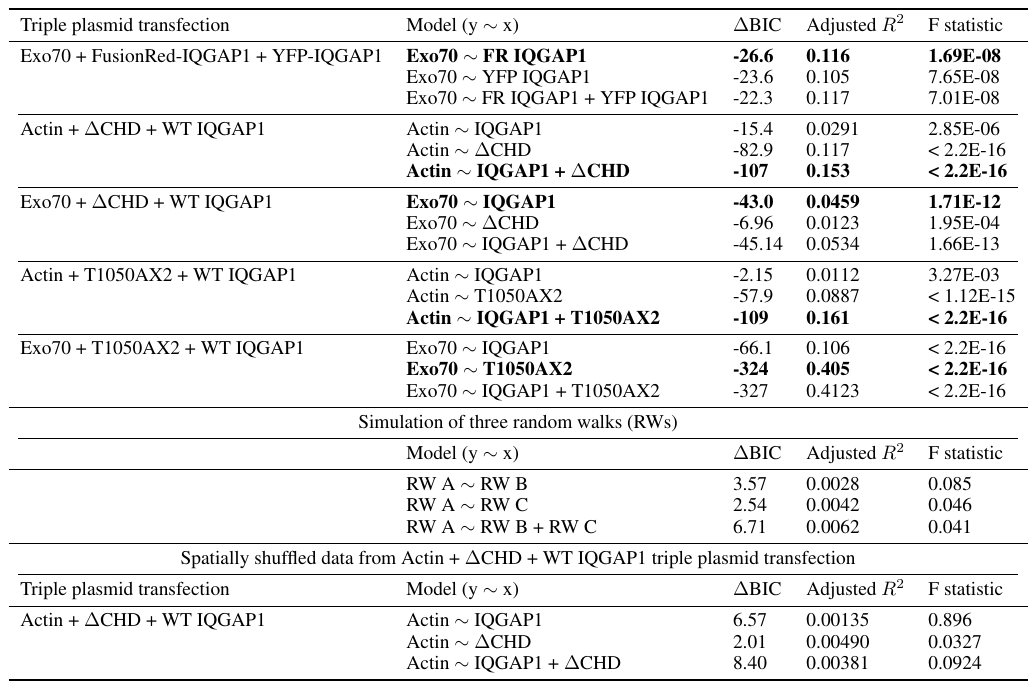

**Control experiments for multiple regression:** Multiple regression methods for local dynamics of scaffold, actin, and membrane signaling components were evaluated in control experiments and simulations.

The first control experiment shows that the choice of fluorescent protein tag does not influence the multiple regression model fit. We transfected MCF10As with a mixture of three plasmids: a WT IQGAP1 construct incorporating an N-terminal FusionRed (FR-IQGAP1), a WT IQGAP1 construct with an N-terminal YFP gene fusion (YFP-IQGAP1), and mTurquoise2-Exo70. Live-cell compartment dynamics were captured with time lapse microscopy and statistically analyzed as before (Materials and Methods). As expected, near-identical ΔBIC and Adjusted R^2^ values were calculated for all linear regression models with Exo70 as the dependent variable (the first row of Table 1 in the main text), suggesting that FR-IQGAP1 dynamics are indistinguishable from YFP-IQGAP1 dynamics. This experiment further demonstrates that using information from two differently-tagged but otherwise identical WT IQGAP1 species does not predict Exo70 dynamics better than a single fluorescent WT IQGAP1 species predicts Exo70 dynamics.

It is known that two random time series often exhibit statistically significant correlation coefficients, and this statistical phenomenon is known as ‘spurious regression’ (Granger and Newbold, 1974). To validate that the fits in our models are not from ‘spurious regression’ we simulated 10,000 triples of random walk (RW) models (analogous to three color triple-protein trajectories) with mean, variance, and number of time points equivalent to those of real compartment trajectories. We then down-sampled each trajectory as we did for the protein trajectories, and constructed regression models analogous to the three models for each triple plasmid transfection. The *p*-values of these models fit to RW simulations are greater by several orders of magnitude than our models of real compartment trajectories (Table S1), verifying that real biophysical events and not spurious regression lead to the observed *p*-values.

A final control analysis verifies our hypothesis that the observed IQGAP1 dynamics are local—specific to single compartments—and not cell-wide. The F-Tractin + ∆CHD + wt-IQGAP1 data set was re-analyzed by computationally shuffling the spatial component (compartment identity) of each intensity measurement. Specifically, while keeping fixed the combination of cell, time point, and fluorescent protein construct, we swapped the intensity measurements from different compartments:
 $I_{\left\{ {cell}_{c}, {time}_{t},{protein}_{p},{compartment}_{i} \right\}}\to I_{\left\{ {cell}_{c}, {time}_{t},{protein}_{p},{compartment}_{j\neq i} \right\}}$.
This procedure created cell-wide, hybrid trajectories, and averaged out the subcellular spatial effects. The resulting low Adjusted R^2^ and positive ∆BIC after spatial shuffling (Table S1) is in stark contrast to the high Adjusted R^2^ and negative ∆BIC before shuffling. This difference suggests the correlative dynamics between measured signaling proteins in this investigation is locally driven at compartments at the micron-scale. For our biological system, this argues against the “well-mixed” approximation to subcellular dynamics that underlies ordinary differential equation models of other cell signaling systems. A further interpretation of the small Adjusted R^2^ after spatial shuffling is that within the time window of our experiments, compartment dynamics is weakly influenced by cell-wide processes such as translation, calcium signaling, and cell cycle events.

Granger, Clive WJ, and Paul Newbold. "Spurious regressions in econometrics." *Journal of Econometrics* 2.2 (1974): 111-120.
